# Behavioural response diversity of individual seabirds can correlate positively with colony stability

**DOI:** 10.64898/2026.11.19.767307

**Authors:** Duncan A. O’Brien, Caitlin K. Frankish, Christopher F. Clements, Pol Capdevila, Hannah S. Wauchope, Annette L. Fayet, Tycho Anker-Nilssen, Manuel Ballesteros, Ingar S. Bringsvor, Nina Dehnhard, Kjell Einar Erikstad, Svein-Håkon Lorentsen, Tone K. Reiertsen, Vegard Sandøy Bråthen, Hallvard Strøm, Geir H. Systad, Arnaud Tarroux, Benjamin Merkel, Kate Layton-Matthews

**Affiliations:** School of Biological Sciences, University of Bristol, Bristol, BS8 1TQ, UK; Norwegian Polar Institute, FRAM Centre, 9296 Tromsø, Norway; Departament de Biologia Evolutiva, Ecologia i Ciències Ambientals, Facultat de Biologia, Universitat de Barcelona (UB), Barcelona, Spain; Institut de Recerca de la Biodiversitat, Universitat de Barcelona (UB), Barcelona, Spain; School of GeoSciences, University of Edinburgh, Edinburgh, UK; Norwegian Institute for Nature Research (NINA), P.O. Box 5685 Torgarden, 7485 Trondheim, Norway; Norwegian Institute for Nature Research (NINA), FRAM Centre, 9296 Tromsø, Norway; BirdLife Møre og Romsdal, Sandshamn, Norway; Norwegian Institute for Nature Research (NINA), 5006 Bergen, Norway; Norwegian Institute for Nature Research (NINA), Oslo, Norway

**Keywords:** conservation, persistence, resilience, theoretical ecology, uncertainty, variance

## Abstract

Ensuring ecosystems are stable has become a priority given humanity’s dependence on the services those ecosystems provide. There has consequently been significant research effort demonstrating that ecosystem stability is driven by ‘response diversity’, whereby different species respond uniquely to the environmental conditions and ‘insure’ the ecosystem against collapse. However, response diversity theory has only been applied at the ecosystem/community level, when the response diversity of individual animals’ behaviour or traits is also theoretically linked to population-level stability. Therefore, we may be overlooking a viable indicator of extinction risk. Using long-term behavioral and energetic expenditure data from two seabird species breeding in the North Atlantic (Atlantic puffin *Fratercula arctica* and black-legged kittiwake *Rissa tridactyla*), we demonstrate that the response diversity in foraging effort of individual birds correlates with the stability of their colony’s population size, while energetic expenditure does not. Further, the diversity of bird responses increases when environmental conditions (here represented by sea surface temperature) are outside of the colony’s norm. Our work shows that response diversity provides a new perspective on the role of individual behavioral differences in creating stable populations, offering insights to better manage vulnerable groups.

## Introduction

Global species extinctions are accelerating [1,2] with demonstrable impacts on the functioning of ecological communities and the resulting services to people [3,4]. Maintaining stable ecosystems is consequently a desirable management outcome and understanding how the diversity of its fundamental units (i.e. species [5]) results in stability has been a key goal of ecological research [6–8].

Ecological stability represents a multidimensional ecosystem characteristic encompassing the ability to both remain unchanged when perturbed and return to equilibrium [6,9]. Theoretical work assessing its relationship with biodiversity has established that higher species richness stabilises community level processes in complex ecosystems [10–12]. The hypothesised mechanism behind this relationship is response diversity [13], or the variation in species responses to environmental change. Specifically, an ecological community supporting a wider range of possible responses to the environment is expected to display asynchrony in its species’ abundances over time [14], a key stabilising mechanism according to the insurance hypothesis [14,15], and a portfolio effect (i.e many species responding uniquely results in statistical averaging at the community level [10,16,17]). *Ergo*, as response diversity drives asynchrony [18,19], and asynchrony drives community stability [15], response diversity is an ultimate cause of stability [20], allowing ecosystems to remain unchanged despite changing conditions [14].

Significant effort has linked species-level response diversity with community stability [18–22], but few attempts have linked intraspecific diversity of individual responses (i.e. among-individual variability) to a species’ local population-level stability in abundance or biomass (e.g. [23]).

Conceptually, response diversity-stability relationships should also be applicable to lower hierarchical levels [24,25] as individuals differ in their phenotypic plasticity (i.e. within-individual variability [26,27]), personalities [28,29] and genotypes [30] to create a range of possible responses to their environment [31]. With different individuals adopting different strategies and, ultimately, fitness outcomes, the potential for a species’ population to increase its ‘population mean fitness’ [32] and buffer changing conditions emerges when a greater range of strategies is present [26,31,33]. However, datasets and model systems capable of exploring response diversity questions of individual organisms remain limited given the challenges of tracking multiple individuals through time [34].

Seabirds represent some of the few intensively monitored organisms able to fill this knowledge gap. Long-term monitoring programmes have measured individual seabird morphology, behaviour and survival across decades [35], providing evidence to define marine protected areas [36], inform fishery regulations [37], and identify drivers of population decline [38].

Previous work has used these datasets to explore how birds from within the same colony vary in migration strategy [39–41], breeding phenology [42], or foraging decisions [27,43,44], demonstrating strong among-individual variation. However, it is unclear whether this individual variation results in stable population-level dynamics. For instance, among-individual variation can correlate with the demographic rates (e.g breeding success and survival) that create stable populations in some seabird systems [28,40,44,45], but not in others [27,41,42,46]. This uncertainty has challenged the usefulness of seabirds as ecosystem indicators [47], as a populations’ relationship with its environmental conditions changes depending on ‘switches’ in plasticity [47]. Reconciling the relationship between individual-level variation and population stability will therefore help define not only risk of population extirpation [26] but also the conditions where ecological indicators can be trusted [48].

Here, we apply response diversity theory to individual level daily behavioral and energetic expenditure data from two North-Atlantic seabird species (Atlantic puffin - *Fratercula arctica* - and black-legged kittiwake - *Rissa tridactyla*), and estimate the relationship between response diversity and population-level stability. Using global location sensor-derived foraging effort as a measure of individual responses to environmental conditions [49,50], and daily energy expenditure as the consequence of those individual foraging decisions [51], we describe how response diversity, measured by previously established dissimilarity (i.e. the difference in response magnitudes) [20] and imbalance (i.e. how centered around zero the average response is) [21] metrics, varies across a sea surface temperature gradient. Finally, we correlate population-level stability, quantified by temporal stability [6] and dynamic eigenvalue metrics [52], with individual-level response diversity, hypothesising that stability positively correlates with among-individual response dissimilarity and negatively correlates with response imbalance.

## Methods

### Seabird and sea surface temperature data

Seabird measurements have been collected across Norway by the national seabird monitoring programme SEAPOP (www.seapop.no) and tracking project SEATRACK (https://seatrack.net). SEAPOP has developed an annual sampling methodology adapted from international standard methodologies (Table S1), which targets breeding adults as those individuals are easier to observe and/or catch as they are associated with their nests. We focussed on two species - Atlantic puffins (*Fratercula arctica*, hereafter puffins) and black-legged kittiwakes (*Rissa tridactyla*, hereafter kittiwakes) - at 6 breeding colonies (i.e. populations, Figure S1). These species likely vary in their sea surface temperature (SST) reaction norms given their differing foraging ecologies (puffins - pursuit divers [53]; kittiwakes - surface-foragers [54]), prey preferences (e.g. puffins - 20-100mm capelin, sandeel; kittiwakes - 100-150mm capelin [55]) and generation time (puffins - 14.2 years; kittiwakes - 9.8 years [56]), and consequently may display unique response diversity relationships with population-level stability. For these species, we estimated each colony’s population size from the sum of apparently occupied nests across fixed sampling plots (Figure S2), accounting for varying sampling effort using TRIM ([57–59] - Supplementary Methods).

Bird behaviour during the breeding season was derived from combined geolocator-immersion loggers (GLS). GLS deployments were coordinated through SEATRACK, which also processed and curated the data (Table S1). These sensors measure light level and conductivity. Raw positions were extracted from the light data and cleaned following Bråthen et al. [60], with missing locations interpolated based on the Informed Random Movement Algorithm (IRMA - [61]). Conductivity data were standardized over 10-minute intervals between 0 (entirely dry) and 1 (entirely wet). Processed light and conductivity data were then time-matched, retaining only individuals with both types of biologging data for subsequent analysis.

We further subset the annual cycle to focus on the breeding season, of which start and end dates were assigned following Fauchald et al. [61] using the 5-day rolling mean across individual birds’ GLS conductivity time series in combination with the Lavielle partitioning algorithm [62]. In brief, this process classifies periods of mostly wet and periods of mostly dry, with the dry periods assumed to represent colony attendance and therefore the breeding season.

We used GLS positions to estimate breeding season utilization kernels for each population (Figure S1) using the adehabitatHR v.0.4.22 [63] R [64] package. As breeding puffins and kittiwakes are central place foragers [53,54], at-sea locations during this period were assumed to represent foraging trips. We then defined each population’s core area as the 50% utilization distribution to mitigate variability due to algorithmic/bandwidth choices. Due to low sample sizes for some population-year combinations, we estimated a single kernel for each colony by pooling individual tracks across all years. From the area defined by each colony’s utilization kernel, we extracted daily sea surface temperatures (SST - °C) from the ECMWF Reanalysis v5 (ERA5) [65], and averaged across each colony’s kernel, assuming all birds experienced these environmental conditions. ERA5 derived daily air temperature (°C 2m above surface) was also extracted at each colony’s coordinates.

Finally, four behaviours; ‘foraging’, ‘resting on water’, ‘resting on land’ or ‘in-flight’ (Figures S3-S4) were derived at a daily temporal resolution from processed light and conductivity data following Christensen-Dalsgaard et al. [50], Fayet et al. [49] and Frankish et al. [66]. These behaviours were assumed to be plastic, with individuals able to make daily decisions [43,49,50]. Foraging effort was then taken forward as an individual-level performance metric appropriate for response diversity calculation. Daily energetic expenditure (DEE) was then estimated from the combination of behavioral classifications and environmental data as described in Frankish et al. ([66] - Supplementary Methods) to represent the ‘consequence’ of within- and among-individual variability in foraging decisions [51]. Due to uncertainty in the parameters used to calculate DEE, we repeated the calculation 1000 times to create a distribution of DEE values.

### Response diversity calculation

To calculate response diversity in seabird daily foraging effort and energetic expenditure, we applied the framework proposed by Ross et al. [20]. Specifically, we used Bayesian generalised additive models (GAMs) to fit performance–environment curves (i.e. reaction norms - Figure 1A) for each colony and species via brms v.2.23.0 [67], allowing each individual bird’s performance– environment smooth to vary independently around a global SST smooth as a random effect.

**Figure 1.**
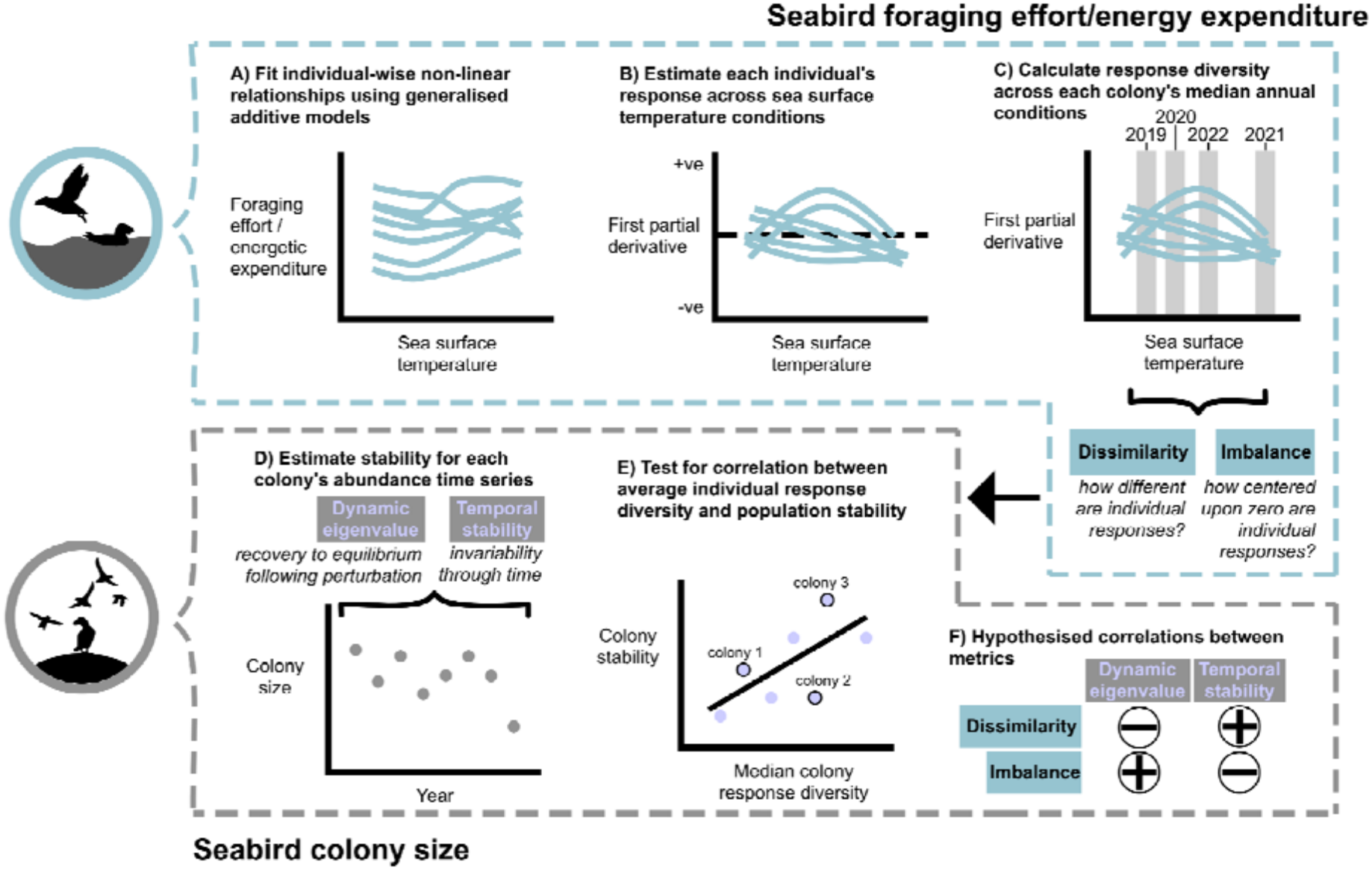
Overview of the analytical procedure for the calculation of response diversity (A-C), population-level stability (D), and the relationship between response diversity and stability (E). Hypothesised correlation direction for each stability-response diversity metric combination is indicated in F.

Sea surface temperature is an established driver of these colonies’ dynamics [35]. Random intercepts for day-of-the-year and year were also included to compensate for varying foraging effort between days/years not explained by SST. The exact model structure fit to the proportional (i.e. [0-1] bounded) foraging effort data was:

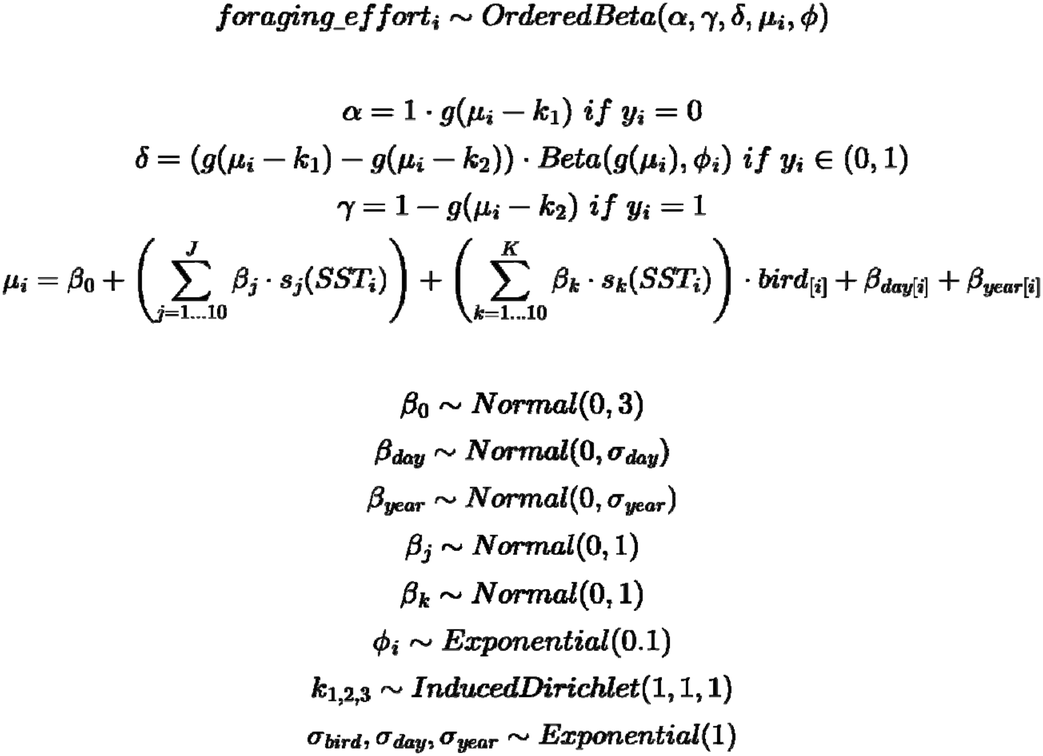

where ***foraging_effort***_*i*_ is the observed foraging effort at index *i*, OrderedBeta is the ordered beta distribution family most appropriate for [0,1] data [68], *α* is the probability of a measurement being 0, *δ* is the probability a measurement is in the continuous range, *γ* is the probability a measurement is 1, and ***k*** is the ordered cutpoints between the components [68].The location linear predictor *µ* was estimated from the coefficients *β* and random errors *σ*, while ***g*(·)** is precision and is the inverse logit function. Smooth functions **(*s*)** are thin plate smooths where the maximum number of knots **(*J*)** was penalised to reduce ***j***. Bird individuals were only modelled if they had been monitored for three or more years.

This model structure was replicated for DEE data, with the ordered beta distribution replaced by a gamma distribution as observations are positive real bounded ([0,∞)), and a measurement error component added:

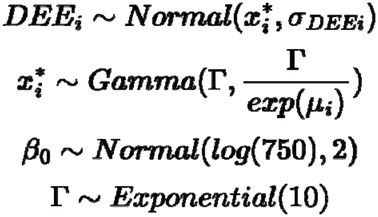

where ***DEE***_*i*_ is the mean estimated DEE at index ***i***, *σ* ***DEE***_*i*_ is the standard deviation of the uncertain DEE estimate, 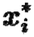 is a latent ‘true’ variable, and Γ is the shape of the gamma distribution. Posterior predictive checks for both foraging effort and DEE GAMs can be found in Figure S5-S6.

Response diversity in foraging effort or DEE was then calculated with uncertainty from the GAM estimated partial derivatives (Figure 1B), via marginaleffects v.0.32.0 [69]. These partial derivatives represent the instantaneous change in foraging effort or DEE for a given SST condition, and are converted to response diversity as either response dissimilarity [20] or imbalance [21] (Figure 1C).

Response dissimilarity is the pairwise Euclidean distance [70] in the partial derivative between all pairs of seabird individuals in the colony for a given SST, with the exponent set to zero [20]. Doing so weighs all individuals as equally important. Lower values of dissimilarity therefore indicate more individuals responding equivalently (in their partial derivatives) to the current SST conditions - i.e low diversity.

Response imbalance was calculated as:

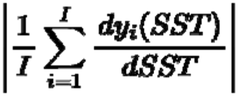

where ***I*** is the number of seabird individuals and 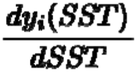 is the instantaneous response of seabird for a given SST (i.e. the first partial derivative) [21]. Response imbalance scales from zero to infinity, where higher values represent lower diversity and all individuals are responding strongly in the same direction (i.e. all positive or all negative). Conversely, low response imbalance indicates high diversity, with ‘net-zero’ average response due to individuals responding weakly to SST, or different individuals responding sufficiently differently from each other such that the average is close to zero [21].

### Stability estimation

Population-level stability was estimated across the entirety of each colony’s detrended population size time series (Figure 1D,F). We focussed on two stability metrics, namely temporal stability [6] and the dynamic eigenvalue (DEV) [52]. Temporal stability (or the invariability of the population through time) was estimated as the inverse of the coefficient of variation, while the DEV uses empirical dynamical modelling to estimate the Jacobian matrix and resulting ‘asymptotic stability’ (i.e. the recovery rate to equilibrium following a perturbation to the colony). High temporal stability and low DEV are both indicative of stability. We used the univariate form of DEV, whereby the colony’s state-space (and Jacobian matrix) is reconstructed via the embedding of the time series against lagged versions of itself [52,71].

Following Grziwotz et al. [52], we set the embedding dimension *E* to 1 (1 year) and the time lag *τ* to 10 *%* the length of the time series. Both stability metrics are reliable estimates of population-level stability [6,9,52,71] and were calculated using EWSmethods v.1.3.3 [72].

### Individual-level response diversity-stability relationships

To determine whether individual response diversity correlates with population-level stability, we summarised response diversity metrics for each year at the median SST each colony experienced across its utilization kernel (Figure 1C). We then calculated the Pearson correlation between overall colony stability (kittiwake: n = 6, puffin: n = 5) with median response diversity across the same time window (Figure 1E) for each species. To propagate uncertainty, we repeated the process for each draw from the response diversity uncertainties to generate a distribution of correlation coefficients.

## Results

In general, foraging effort was higher and more variable in kittiwakes compared to puffins (kittiwake mean foraging effort [SD] = 0.29 proportion of daylight hours [0.17], puffins = 0.20 proportion of daylight hours [0.14]), while the opposite was true for daily energetic expenditure (DEE) (kittiwakes = 638.59 kJ [249.02], puffins = 796.45 kJ [223.38]). Both species displayed non-linear reaction norms in relation to sea surface temperature (SST) that varied between species and colonies (Figures 2, S7-S8).

**Figure 2.**
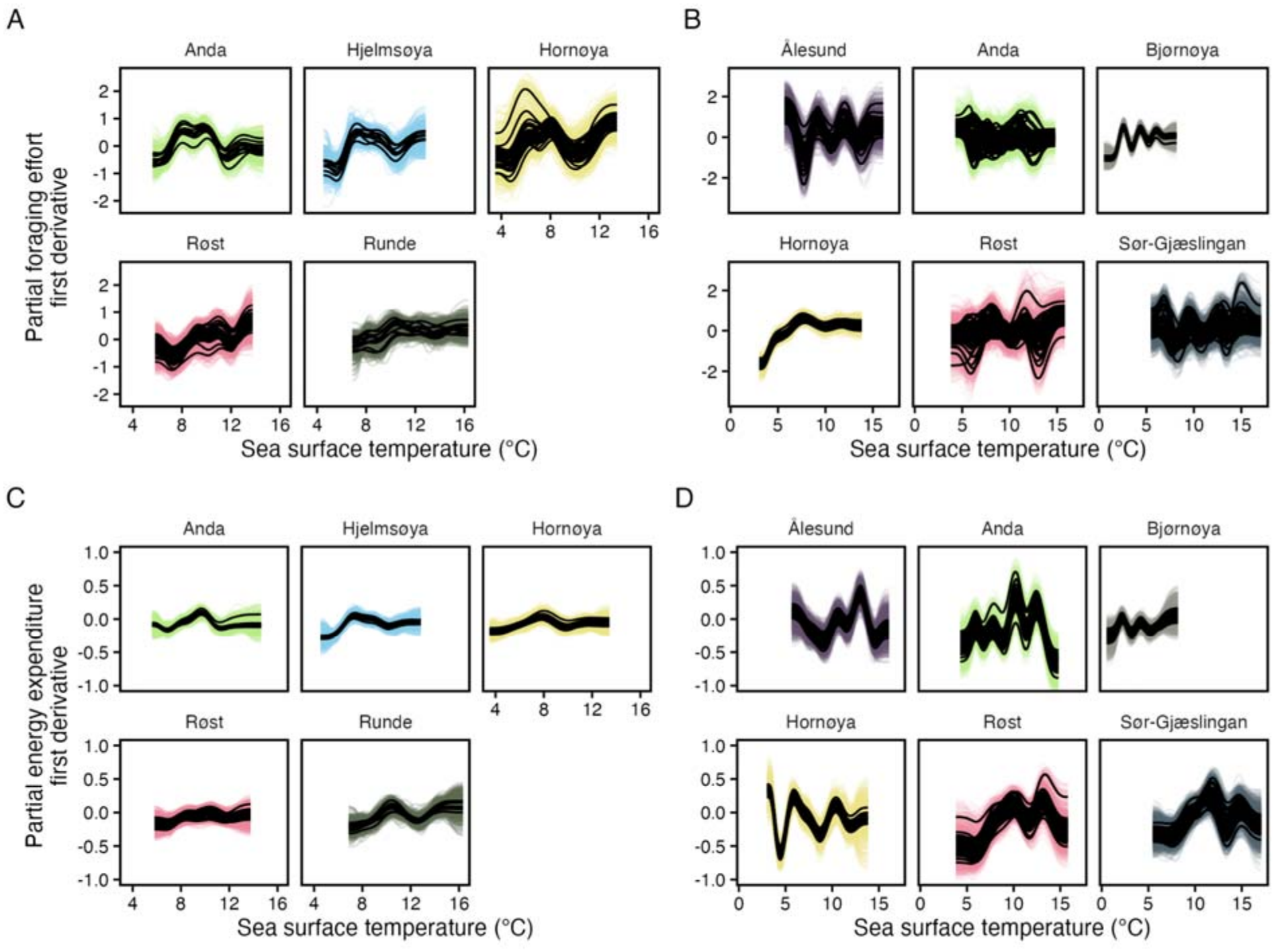
Estimated first partial derivatives of A) puffin and B) kittiwake foraging effort reaction norms across sea surface temperature, and C) puffin and D) kittiwake daily energetic expenditure reaction norms by colony. Solid, dark lines are each individual bird’s median derivative, while coloured lines are 50 random draws for each. These derivatives represent the ‘raw’ data from which response diversity is calculated.

### Response diversity is higher when environmental conditions are atypical

Kittiwakes and puffins displayed similar foraging effort response dissimilarity and imbalance (kittiwake median dissimilarity [95% credible interval] = 1.52 [1.09, 1.98], puffin dissimilarity = 1.27 [1.15, 1.56], kittiwake imbalance = 0.26 [0.01, 1.21], puffin imbalance = 0.32 [0.02, 0.90]), whereby higher response dissimilarity and lower response imbalance are both indicative of higher response diversity. Response dissimilarity specifically expressed a U shape along the SST gradient (Figures S8-S9), with a greater diversity of bird behaviours away from the colony/population’s average SST conditions. This relationship was repeated for kittiwake response imbalance but did not hold in puffins (Figures S9-S10), where a W shape often emerged. Conversely, DEE response dissimilarity displayed no consistent relationship across species nor colonies (Figures S11-S12), though response imbalance was generally U-shaped. Overall DEE response diversity was higher in kittiwakes than puffins (kittiwake dissimilarity = 1.10 [1.05, 1.18], puffin dissimilarity = 1.04 [1.02, 1.07], kittiwake imbalance = 0.16 [0.01, 0.57], puffin imbalance = 0.07 [0.00, 0.25], Figures S10-S11). Kittiwakes from the higher latitude colonies (Bjørnøya and Hornøya) were atypical relative to other colonies in their foraging effort and DEE response diversity, with birds responding similarly regardless of the SST conditions.

### Behavioral response diversity often correlates with stability

Corresponding to our hypotheses, average kittiwake foraging effort response diversity consistently correlated with population-level stability (Figure 3A, Table S2). Kittiwake response dissimilarity was negatively correlated with the dominant eigenvalue (DEV) (median rho [95% credible interval] = −0.87 [−0.94, −0.77]) while response imbalance was positively correlated (0.69 [0.30, 0.66]). Equivalently, kittiwake response dissimilarity was positively correlated with temporal stability (0.49 [0.30, 0.66]) while response imbalance was negatively correlated (−0.38 [−0.77, −0.08]). However, in puffins, average foraging effort response diversity only correlated with DEV: puffin DEV negatively correlated with response dissimilarity (−0.73 [−0.88, −0.50]). The remainder of correlations were either zero or in the opposite to the hypothesised direction (Figure 3A, Table S2).

**Figure 3.**
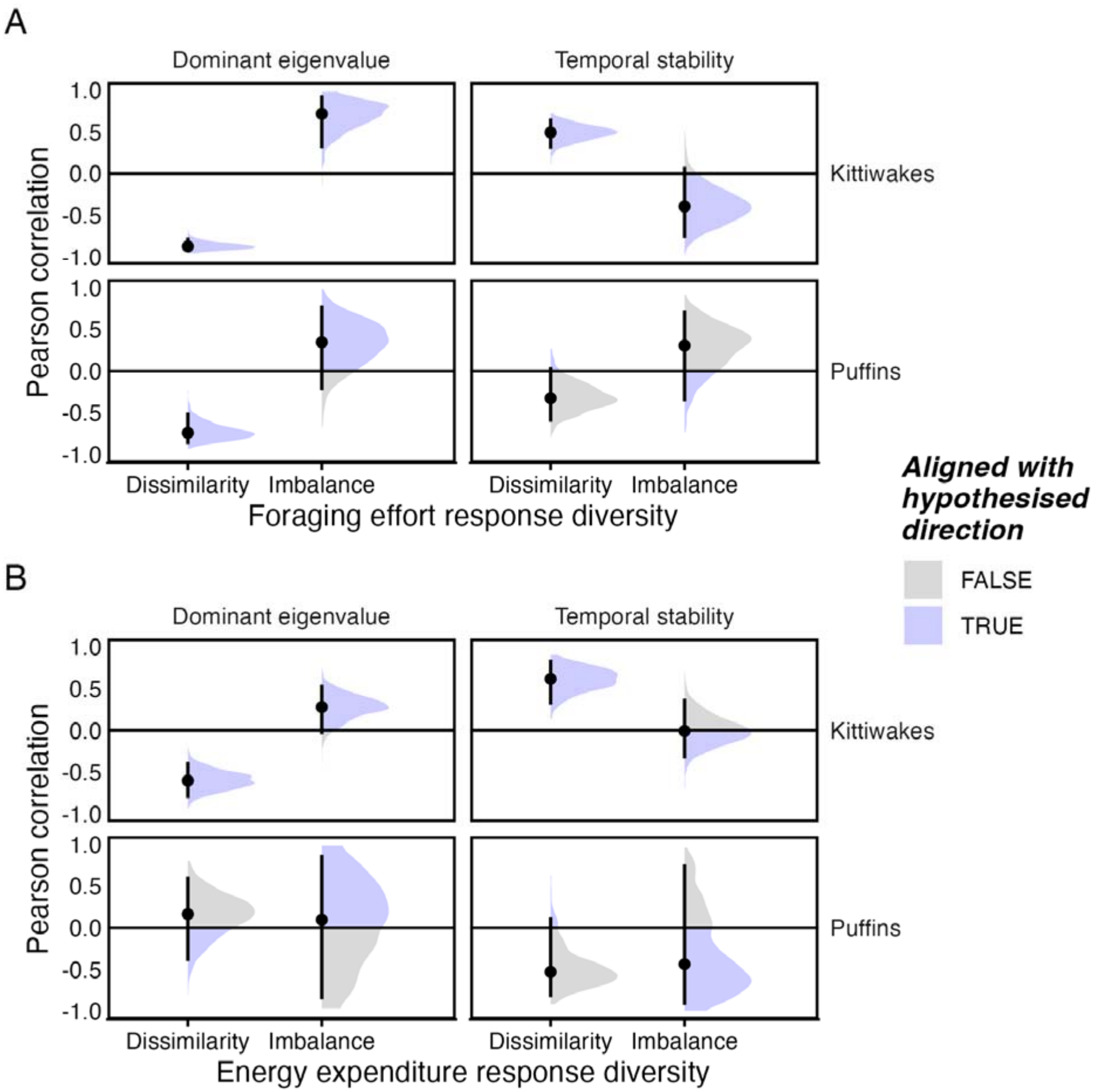
Pearson correlation coefficient between seabird population stability (as represented by the dominant eigenvalue and temporal stability) and A) foraging effort and B) daily energetic expenditure response diversity (represented by dissimilarity and imbalance). Each density plot represents 1000 samples from the posterior distribution of the slope estimate, with median values (circles) and 95% credible intervals (bars) indicated.

Average colony DEE response diversity showed few associations with either measure of population stability: only kittiwake response dissimilarity negatively correlated with DEV (Figure 3B, Table S3; −0.60 [−0.81, −0.38]) and positively with temporal stability (0.61 [0.31, 0.84]).

## Discussion

Understanding why animal populations persist is an active area of research interest [26,34]. Among-individual variation and, more broadly, response diversity has emerged as one lens to interrogate these population level questions, but only once has it been directly compared to population stability [23]. We consequently applied theory and metrics developed in community ecology [10,15,20,21] to individual seabird data, finding foraging effort response diversity increased when colonies deviated from their typical sea surface temperature (SST) conditions. In addition, foraging effort response diversity correlated with population stability as hypothesised, but we detected no relationship between stability and daily energy expenditure (DEE) variation.

Our observation that puffin and kittiwake individuals differed most in their foraging effort responses while away from their typical conditions matches new relationships observed between species asynchrony and stability [73]. It also supports Grémillet & Charmantier’s [47] argument that seabirds’ phenotypic plasticity complicates their use as marine indicators. This raises interesting evolutionary questions around why individuals differ more when conditions are poor (i.e. at high or low SST) and it is, potentially, the ‘population plasticity’ (i.e. the among-individual variation) itself that is informative as an indicator. Similarly, the ecology of a species is likely to influence both their value as an indicator and the strength of any response diversity-stability relationships. For instance, we observed clear kittiwake behavioral response diversity relationships matching our hypothesis, but puffins were inconsistent. Broadly, puffins are more capable of buffering stressful conditions given their longer generation times [56] and an ability to access otherwise inaccessible prey resources by diving [53]. This would erode the relationship between response diversity and stability as time delays will emerge due to recruitment effects [74], and/or individuals will display lower variability in behaviour, as we observed. That said, our findings suggest that loss of individual foraging effort response diversity can indicate population instability and so we encourage further work developing individual-level response diversity theory for conservation/management purposes.

Specifically, while response diversity has the potential to identify how among-individual variability underpins buffering by seabird populations, we do not yet know how the resulting asynchrony/statistical averaging [10,15] alters demographic rates and ultimately population-level stability. This is demonstrated by the observed species-specific diversity-stability relationships and conflicting evidence for the effect of seabird among-individual variation upon said rates and individual fitness [27,28,40,41,46]. Although this work represents the first steps towards understanding the scaling up of individual bird variability to the population level, it is based upon a small sample of populations and individuals (~1% of apparently occupied nests) due to the challenge of collecting appropriate data [34]. For example, of the populations monitored here, most have experienced some degree of decline (Figure S2), decreasing the range of possible stabilities to correlate across. We also do not compensate for other forms of plasticity such as social learning which has been demonstrated to interact with passerine among-individual variability to buffer novel conditions [33]. It is plausible kittiwakes and puffins display similar processes as both species ‘flock’ to foraging areas based upon the presence of conspecifics [75,76]. These extensions are therefore opportunities for future applications of response diversity to individual-level data.

While foraging effort response diversity trends behaved as expected, the lack of DEE relationships is unsurprising given the coarse estimates involved and uncertainty around the plasticity of DEE [77,78]. Further, we expect that different behavioural responses would translate into different DEE consequences and, ultimately, different fitness outcomes, rather than individuals actively altering metabolic rates. As the DEE calculation only considers half the energy budget, with no information on energy intake, different behavioural strategies (as represented by foraging effort for example) can result in similar DEE estimates given the same environmental conditions [51]. Alternatively, physiological constraints may exist that limit variability of DEE during the breeding season for these species [78]. This therefore implies DEE provides less value than direct behavioral estimates within the population-stability framework.

To conclude, individual-level response diversity offers a promising avenue for contextualizing the stability of animal populations. There is a growing need to forecast the future of imperilled animals with non-abundance measurements, such as genotype, behavior or morphology, representing plausible early indicators of decline [26]. We therefore suggest that response diversity provides a framework for linking behavioral ecology within broader ecological processes.

## Data availability statement

Raw tracking data is available following a data request from SEATRACK (https://seatrack.net) whereas SEAPOP abundance data is available from the Norwegian Institute for Nature Research data portal (https://www2.nina.no/seapop/seapophtml/). ERA5 atmospheric data (air and sea surface temperature) can be found through the Copernicus Climate Change Service Climate Data Store (https://doi.org/10.24381/cds.4991cf48). All code and models are available in the public repository: https://github.com/duncanobrien/seabird-RD.

## Acknowledgments

This study was supported by UKRI grants NE/Z000130/1, UKRI1287 and UKRI4325. Data collected through the SEAPOP programme (www.seapop.no/en) was financed by the Norwegian Environment Agency, the Norwegian Ministry of Energy via the Research Council of Norway (Grant number 192141) and Offshore Norge. The SEATRACK programme (https://seatrack.net) is funded by the Norwegian Ministry of Climate and Environment, the Norwegian Ministry of Energy via the Research Council of Norway, the Norwegian Environment Agency, the Norwegian Coastal Administration and Offshore Norge along with 16 energy companies. Data were also collected by the Norwegian Polar Institute (via the Monitoring of Svalbard and Jan Mayen (MOSJ) program, www.npolar.mosj.no). Data collected prior to SEATRACK and SEAPOP received financial support from the preceding institutions of the Norwegian Environment Agency and Equinor, as well as the Research Council of Norway (Hornøya, Grant number 216547), the Norwegian Polar Institute and the County Governors of Nordland, Troms, and Finnmark. Permits to work in the colonies were obtained from the local County Governors, with permissions to handle birds and equip them with loggers granted by the Norwegian Environment Agency and the Norwegian Food Safety Authority. We thank all those responsible for the seabird tracking and monitoring at the different sites and their numerous field assistants and students who contributed to the data collection over the years. We especially thank Signe Christensen-Dalsgaard for Anda and Runde. Finally, we thank the Norwegian Coastal Administration for facilitating stays at Sklinna and Anda lighthouses, and Hornøyas venner for Hornøya lighthouse.

