## Supplementary Materials for "Behavioural response diversity of individual seabirds can correlate positively with colony stability"

#### Contents

**S1: Supplementary methods** **2**

**S2: Supplementary figures** **6**

**S3: Supplementary tables** **18**

**Supplementary references** **22**

### S1: Supplementary methods

##### Estimating population abundance using TRIM (TRends and Indices for Monitoring data - Pannekoek and van Strien 2005)

SEAPOP surveys static sampling plots at each seabird colony for the number of apparently occupied burrows (puffins) or apparently occupied nests (kittiwakes). However, the number of plots sampled within a colony can vary by year due to logistical constraints. We therefore compensated for this varying sampling effort by interpolating each sampling plot that was unsampled in certain years using the generalised TRIM model (Barett et al. 2006, Knape 2016):

$$Var(y_{i,t}) = \theta\mu_{i,t}$$

$$\mu_{i,t} = \beta_0 + \beta_{year[t]} + \beta_{plot[i]}$$

where $y$ is each plot's ($i$) count in a given year ($t$), for a given mean ($\mu$) and dispersion ($theta$). Specifically, we fit a global intercept ($\beta_0$) with year and plot varying fixed effects ($\beta_{year}$,$\beta_{plot}$).

We then predicted expected counts for the missing plots, summing counts across all measured and interpolated plots in a given year to provide an estimate of total apparently occupied burrows/nest per year and colony. Only four colonies required interpolation: Hjelmsøya and Hornøya puffins and Hornøya and Røst kittiwakes.

##### Røst *Rissa tridactyla* abundance

Røst kittiwakes required additional data processing as colony-level population size was measured across two sites (Vedøy and Harbour) which display different trends and consequently different insights into the fate of this colony (i.e. Vedøy supported a large population which declined over 40 years to extinction in 2020, whereas Harbour is a much smaller colony that emerged in 1980, grew, and now is in decline - Anker-Nilssen et al. 2023). While maximum apparently occupied nests were similar between the two sites, Vedøy counts represented 2% sampling effort (estimated from a true versus sampling plot count comparison made in 1988) whereas it was 100% at Harbour. We therefore estimated Røst kittiwake population size as the weighted average between the two sites using the following equation:

$$y_{røst} = \frac{w_{vedøy} \cdot y_{vedøy} + w_{harbour} \cdot y_{harbour}}{w_{vedøy} + w_{harbour}}$$

$$w_{vedøy} = \frac{\Lambda \cdot nplots_{vedøy}}{\Lambda \cdot nplots_{vedøy} + nplots_{harbour}}$$

$$w_{harbour} = \frac{nplots_{harbour}}{\Lambda \cdot nplots_{vedøy} + nplots_{harbour}}$$

$$\Lambda = \frac{y_{true}}{y_{sample}}$$

where $w$ is the site's weight, $y$ is the site's population size summed across all sampling plots, $nplots$ is the number of sampling plots at the site, and $\Lambda$ is the ratio of true kittiwake count to total sample plot count on Vedøy in 1988.

#### ***Fractercula artica* daily energetic expenditure (Frankish et al. 2026)**

Puffin daily energetic expenditure (DEE) was estimated from equations parameterised by Brünnich's guillemot activity-specific cost estimates (Buckingham et al., 2025; Croll et al., 1992; Elliott et al., 2013; Elliott & Gaston, 2014; Gabrielsen et al., 1988; Patterson et al., 2022). As these Brünnich's guillemot costs ($x$) are reported in kJ, we converted them to puffin appropriate costs assuming the following allometric scaling (Shaffer 2011):

$$puffin\ kJ\ g^{-1} = \frac{x\ kJ}{(x\ \overline{g})^{0.689}} \cdot (puffin\ \overline{g})^{0.689}$$

$$x\ \overline{g} = 803$$

$$puffin\ \overline{g} = 395$$

where mean Brünnich's guillemot mass in grams $x(\overline{g})$ is reported in Croll & McLaren (1993), and mean puffin mass is calculated from all puffins captured during this study.

Lower critical temperature in water ($\mathrm{LCT_{water}}$), representing the additional cost of activity incurred in water due to thermoregulation, was derived from experimental estimates of $\mathrm{LCT_{water}}$ in

Brünnich's guillemots (14.18°C - Croll & McLaren 1993; Buckingham et al. 2025). Lower critical temperature in air ($LCT_{air}$) was then derived from Ellis & Gabrielsen (2002 - 5.72°C), assuming a breeding latitude of 78.2°.

Daily energetic expenditure was then estimated from the activity budgets using the following equations:

$$DEE\ kJ\ g^{-1} = 4.35 \cdot T_{flight} + (1.18 - 0.03 \cdot SST) \cdot T_{forage} + (0.31 - 0.005 \cdot TAS) \cdot T_{restland} + (0.72 - 0.03 \cdot SST) \cdot T_{restwater} \quad if\ (\ SST \leq LCT_{water} \wedge TAS \leq LCT_{air})$$

$$DEE\ kJ\ g^{-1} = 4.35 \cdot T_{flight} + 0.79 \cdot T_{forage} + 0.28 \cdot T_{restland} + 0.28 \cdot T_{restwater} \quad if\ (\ SST > LCT_{water} \wedge TAS > LCT_{air})$$

where $T_{activity}$ is hours spent in each activity, $SST$ is sea surface temperature (°C) and $TAS$ is air temperature (°C), assuming that 'foraging' is equivalent to 'active' in Frankish et al. (2026).

##### ***Rissa tridactyla*** **daily energetic expenditure (Frankish et al. 2026)**

Kittiwake daily energetic expenditure (DEE) was estimated using a combination of activity and thermoregulatory costs reported in Gabrielsen et al. (1988), Humphreys et al. (2007) and Tremblay et al. (2024). Specifically, costs were derived from Tremblay et al. (2024), whereby flight costs are represented by 'flapping flight' cost, foraging costs are the mean of 'swimming' and 'flapping flight', and resting-on-water costs the mean of 'swimming' and 'resting'.

All activity cost estimates ($x$) were converted from mL $O^2$ $g^{-1}$ $h^{-1}$ to kJ $g^{-1}$ $h^{-1}$ assuming a caloric conversion factor of 20.1 J $mL^{-1}$ (Schmidt-Nielsen 1997), and rescaled allometrically (Shaffer 2011):

$$kittiwake\ kJ\ g^{-1} = \frac{x\ kJ}{(x\ \overline{g})^{0.717}} \cdot (kittiwake\ \overline{g})^{0.717}$$

$$x\ \overline{g} = 365 \quad if\ Gabrielsen\ et\ al.\ (1998)$$

$$x\ \overline{g} = 450 \quad if\ Tremblay\ et\ al.\ (2024)$$

$$kittiwake\ \overline{g} = 392$$

Lower critical temperature's were revised from Gabrielsen et al. (1988)'s Scholander curve between thermoregulation cost and air temperature, assuming thermal conductivity in water is 2.14 times higher than in air as demonstrated in guillemots (Croll & McLaren 1993 - 4.5°C). Lower critical temperature in water was adjusted 12.5°C to match experimental kittiwake evidence (Humphreys et al. 2007).

Daily energetic expenditure was then estimated from the activity budgets using the following equations:

$$DEE\ kJ\ g^{-1} = 0.91 \cdot T_{flight} + 0.70 \cdot T_{forage} + (0.16 - 0.005 \cdot TAS) \cdot T_{restland} + (0.54 - 0.01 \cdot SST) \cdot T_{restwater} \quad if\ (\ SST \leq LCT_{water} \wedge TAS \leq LCT_{air})$$

$$DEE\ kJ\ g^{-1} = 0.91 \cdot T_{flight} + 0.70 \cdot T_{forage} + 0.13 \cdot T_{restland} + 0.39 \cdot T_{restwater}\ if\ (\ SST > LCT_{water} \wedge TAS > LCT_{air})$$

where $T_{activity}$ is hours spent in each activity, $SST$ is sea surface temperature (°C) and $TAS$ is air temperature (°C).

#### S2: Supplementary figures

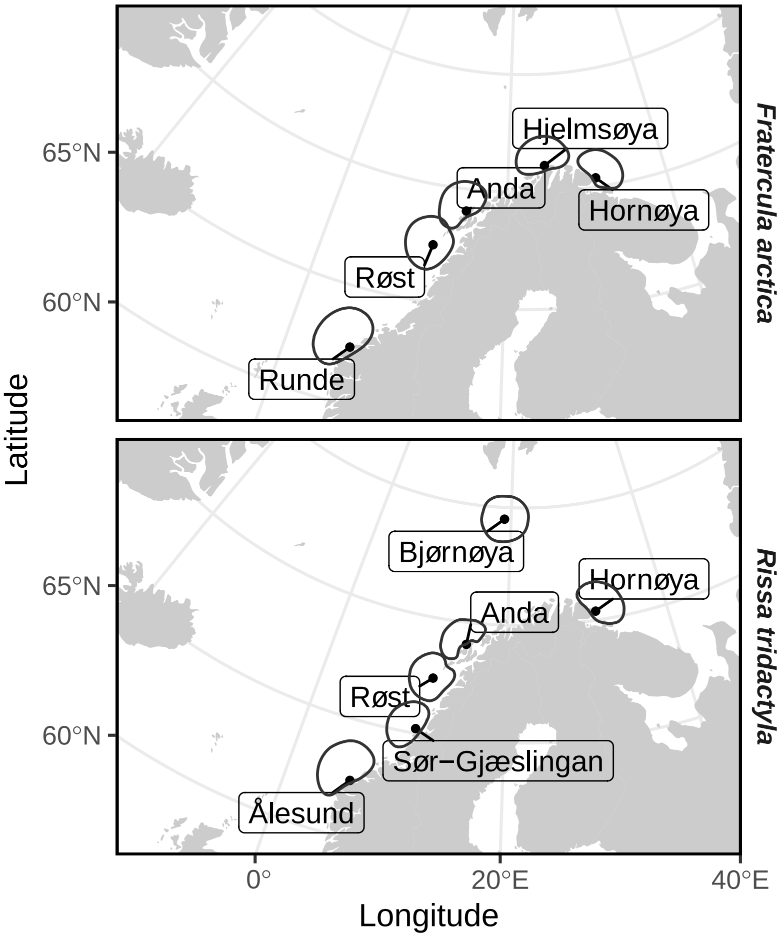


Figure S1: Seabird colony locations (points) and core area kernels (black lined polygons).

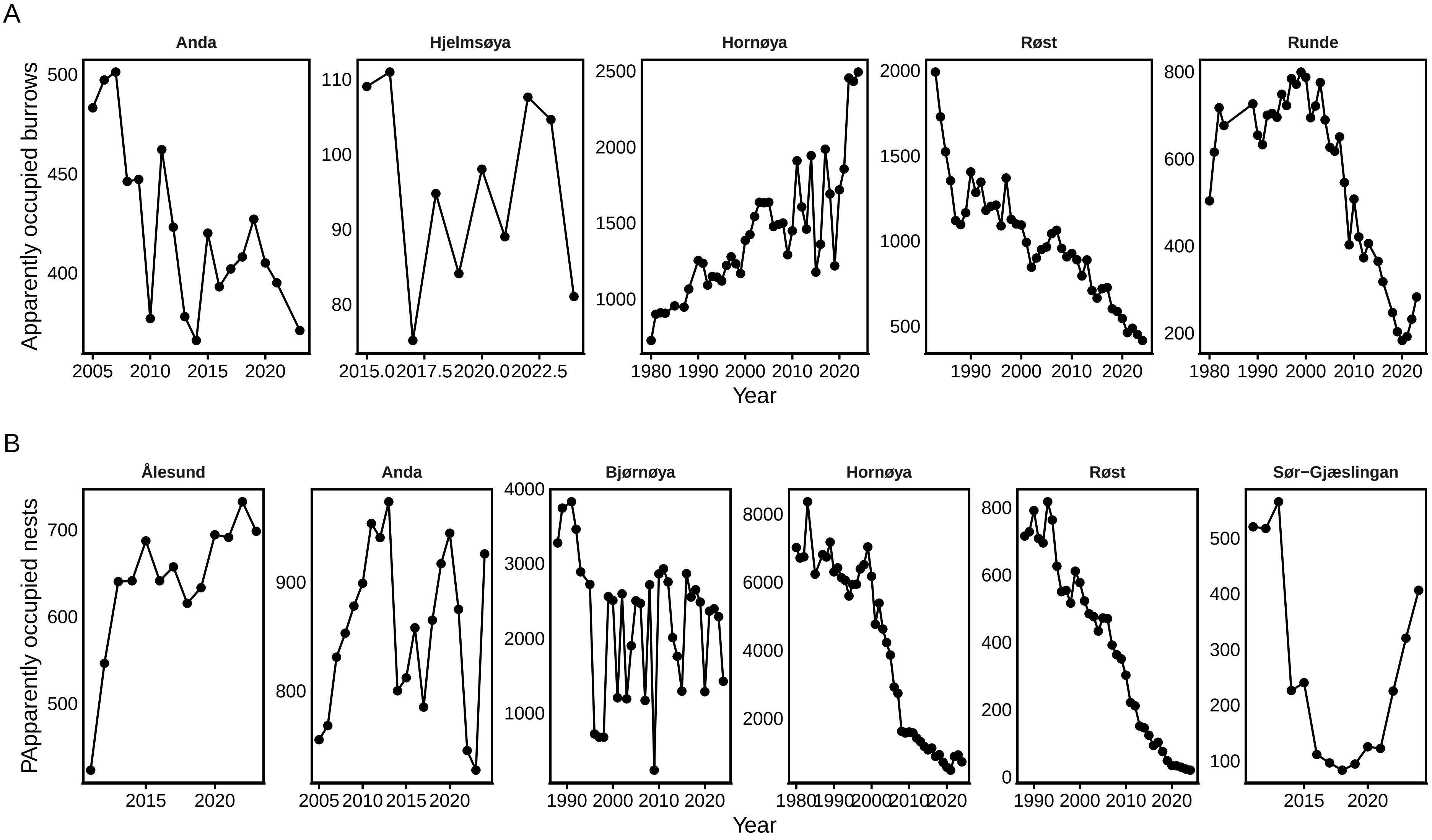


Figure S2: Population size through time in A) *Fratercula arctica* (Atlantic puffin) and B) *Rissa tridactyla* (black-legged kittiwake) colonies.

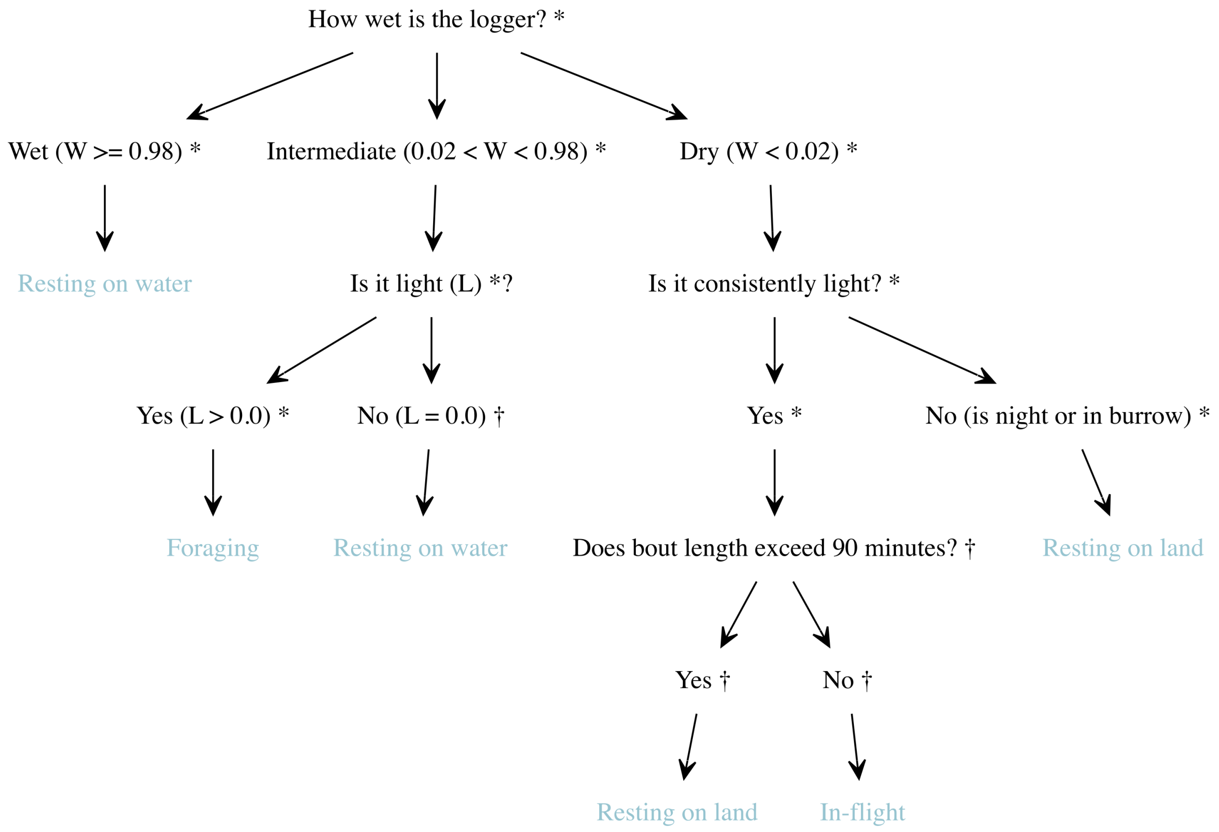


Figure S3: Behavioural decision tree for assigning 10-minute bouts of immersion and light data in *Fratercula arctica* (Atlantic puffin). Data sources: * Fayet et al. 2016, † Frankish et al. 2026.

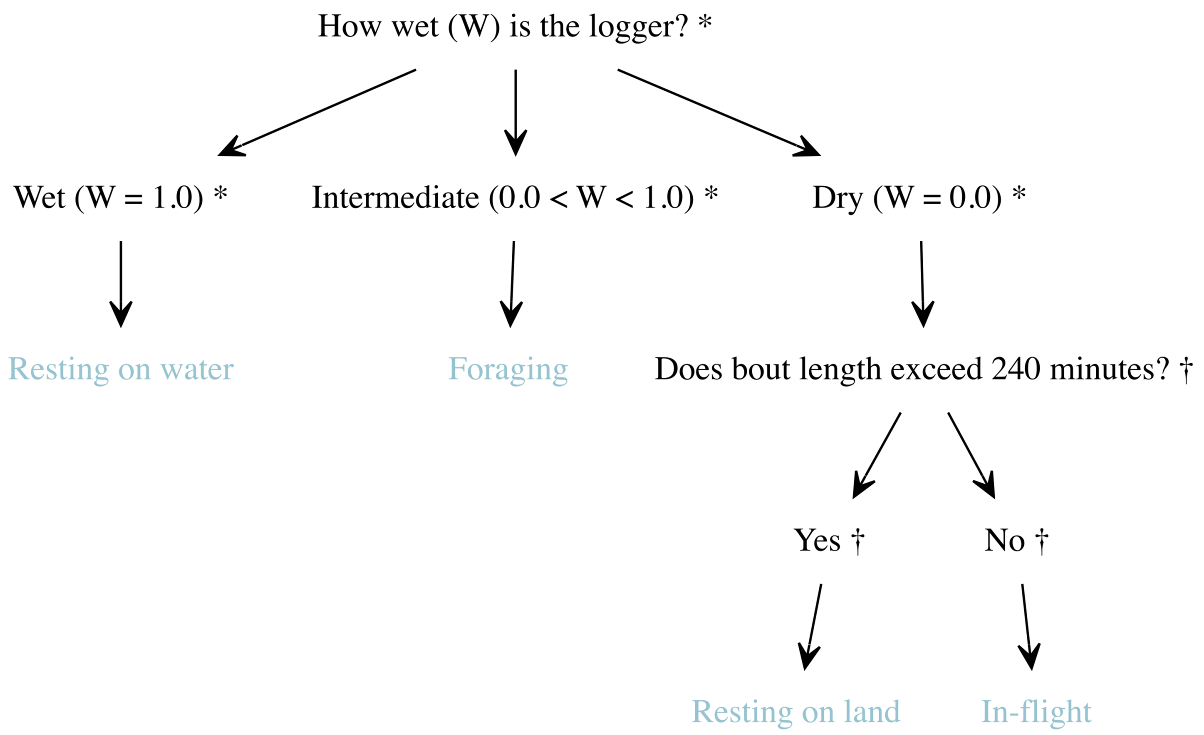


Figure S4: Behavioural decision tree for assigning 10-minute bouts of immersion and light data in *Rissa tridactyla* (black-legged kittiwake). Data sources: * Christensen-Dalsgaard et al. 2018, † Frankish et al. 2026.

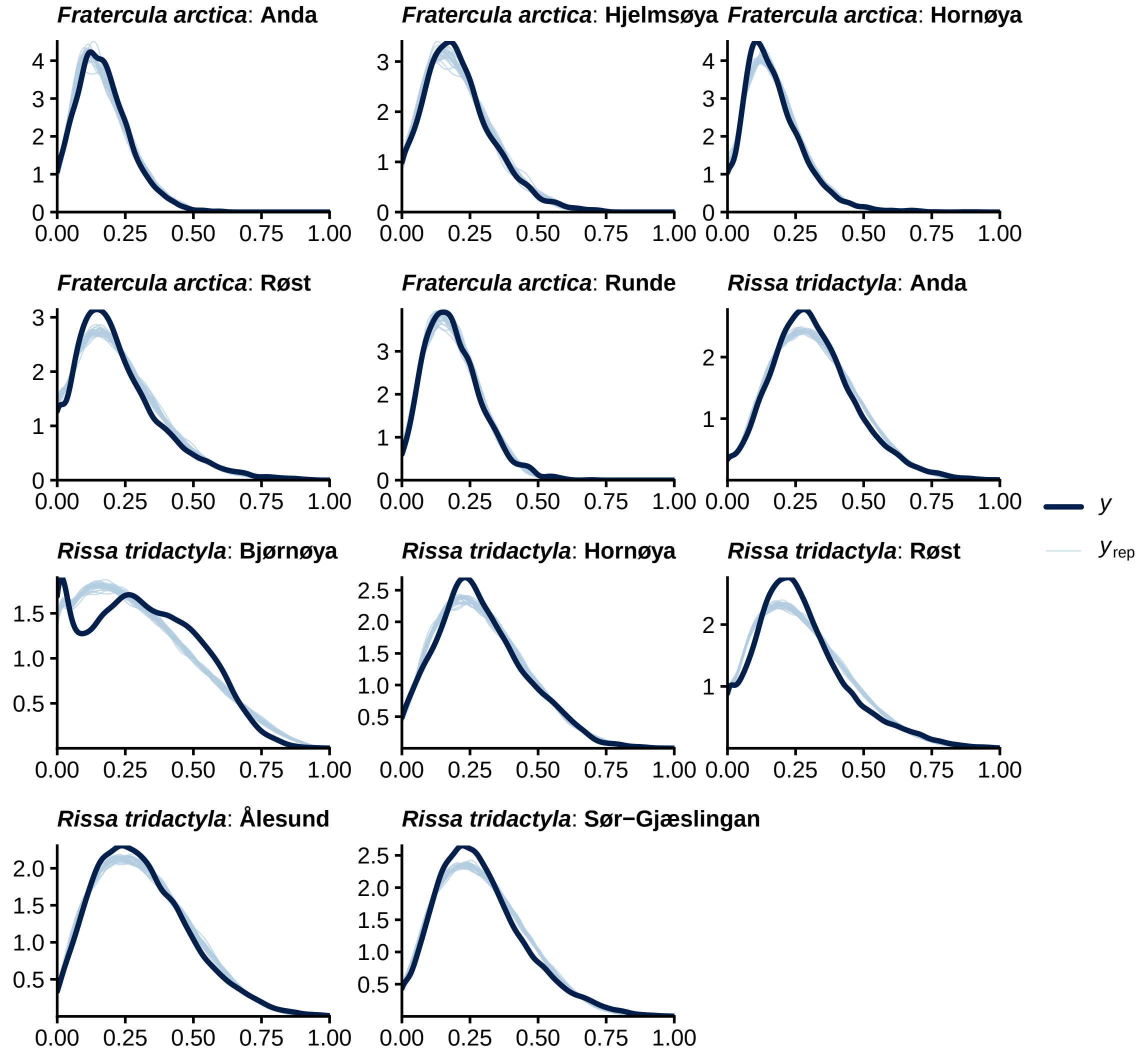


Figure S5: Posterior predictive checks for ordered beta generalised additive models fit to estimate foraging effort reaction norms. Thick dark lines ($y$) are observed data, and thin light lines are ($yrep$) are 25 random draws from the model posterior.

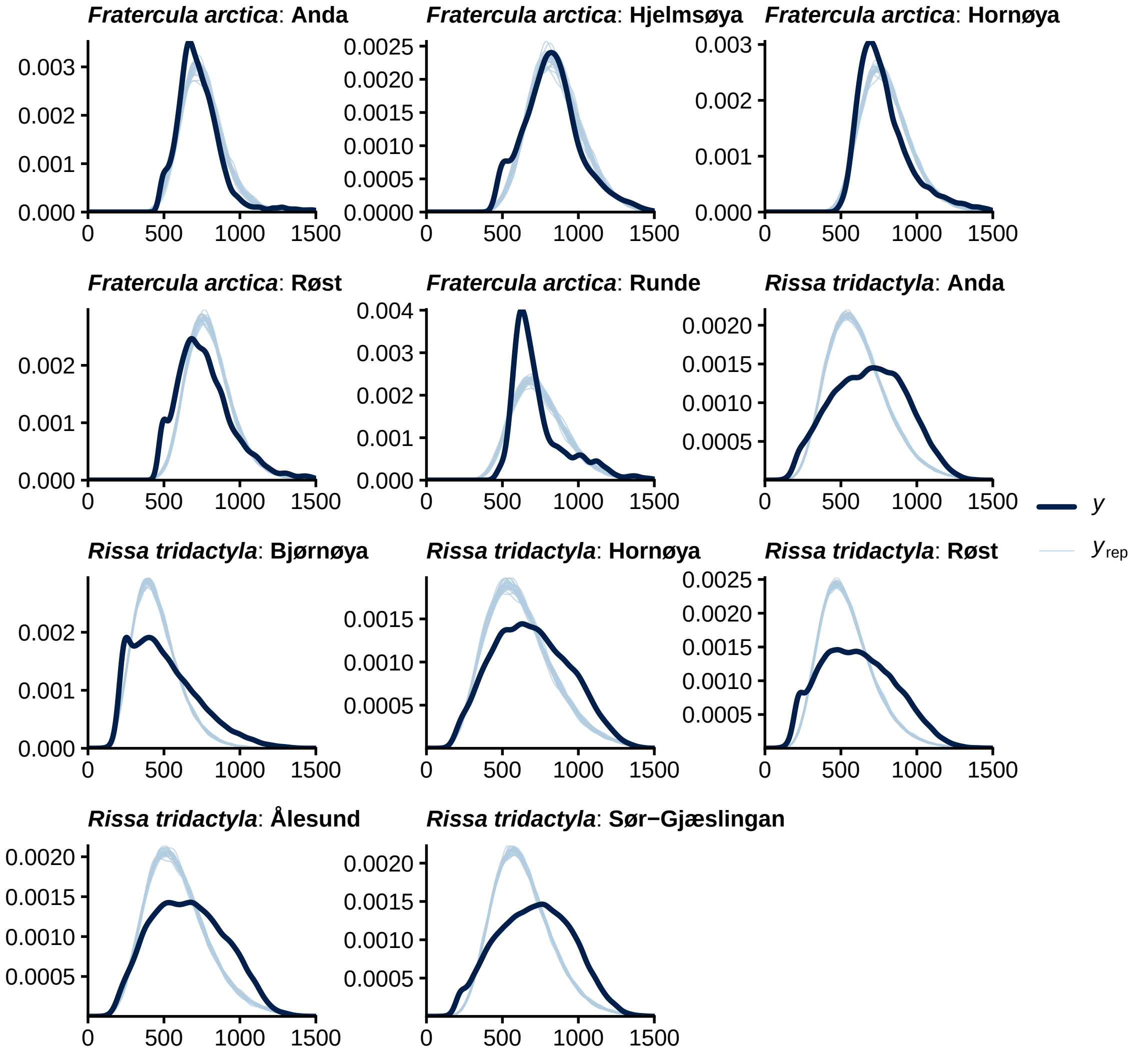


Figure S6: Posterior predictive checks for gamma generalised additive models fit to estimate daily energetic expenditure reaction norms. Thick dark lines ($y$) are observed data, and thin light lines are ($yrep$) are 25 random draws from the model posterior.

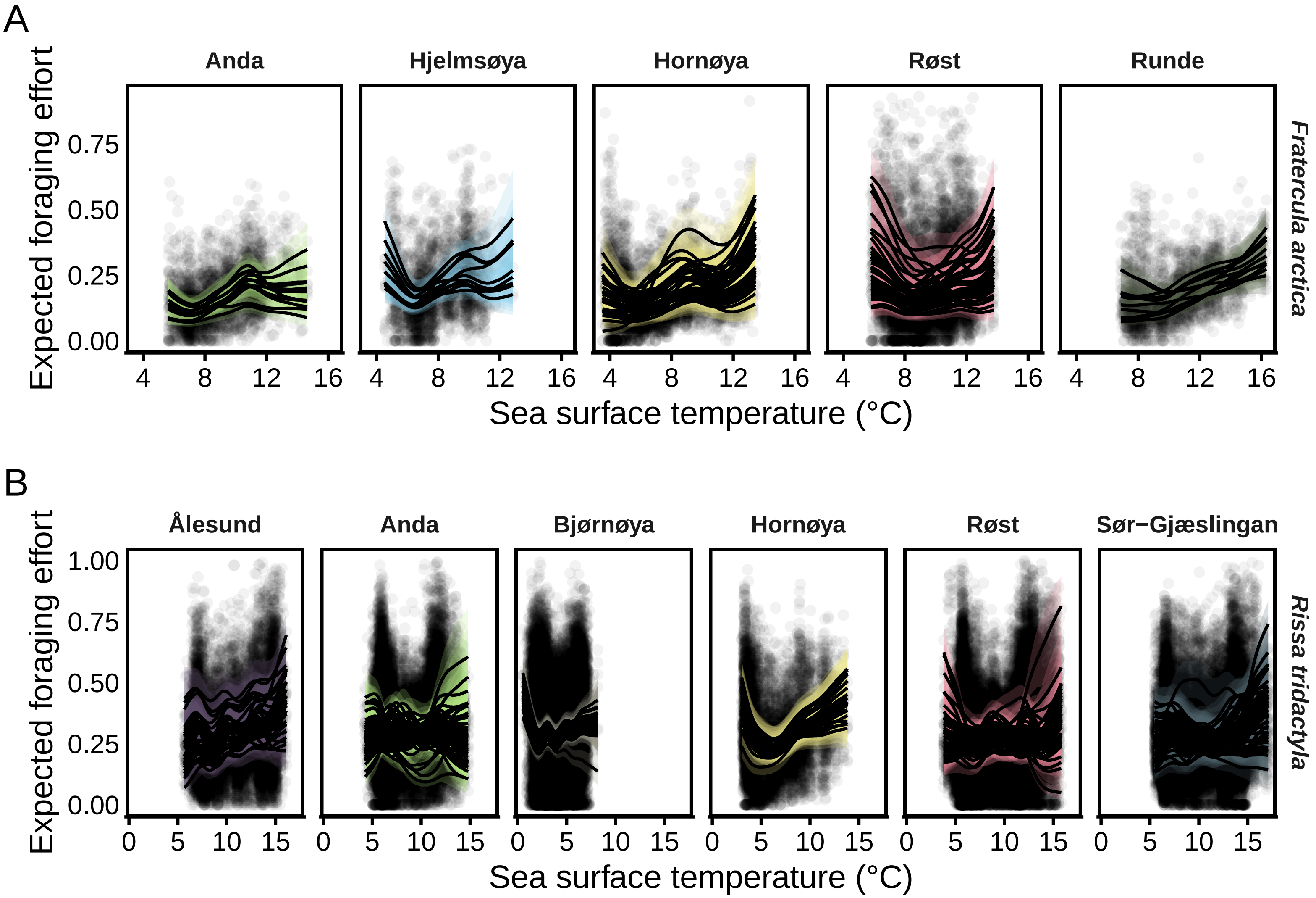


Figure S7: Expected foraging effort (proportion of day spent foraging) for each individual and colony, in A) puffins and B) kittiwakes, as predicted by generalised additive models. Solid lines are median estimates, with shaded regions the pointwise 95% credible interval.

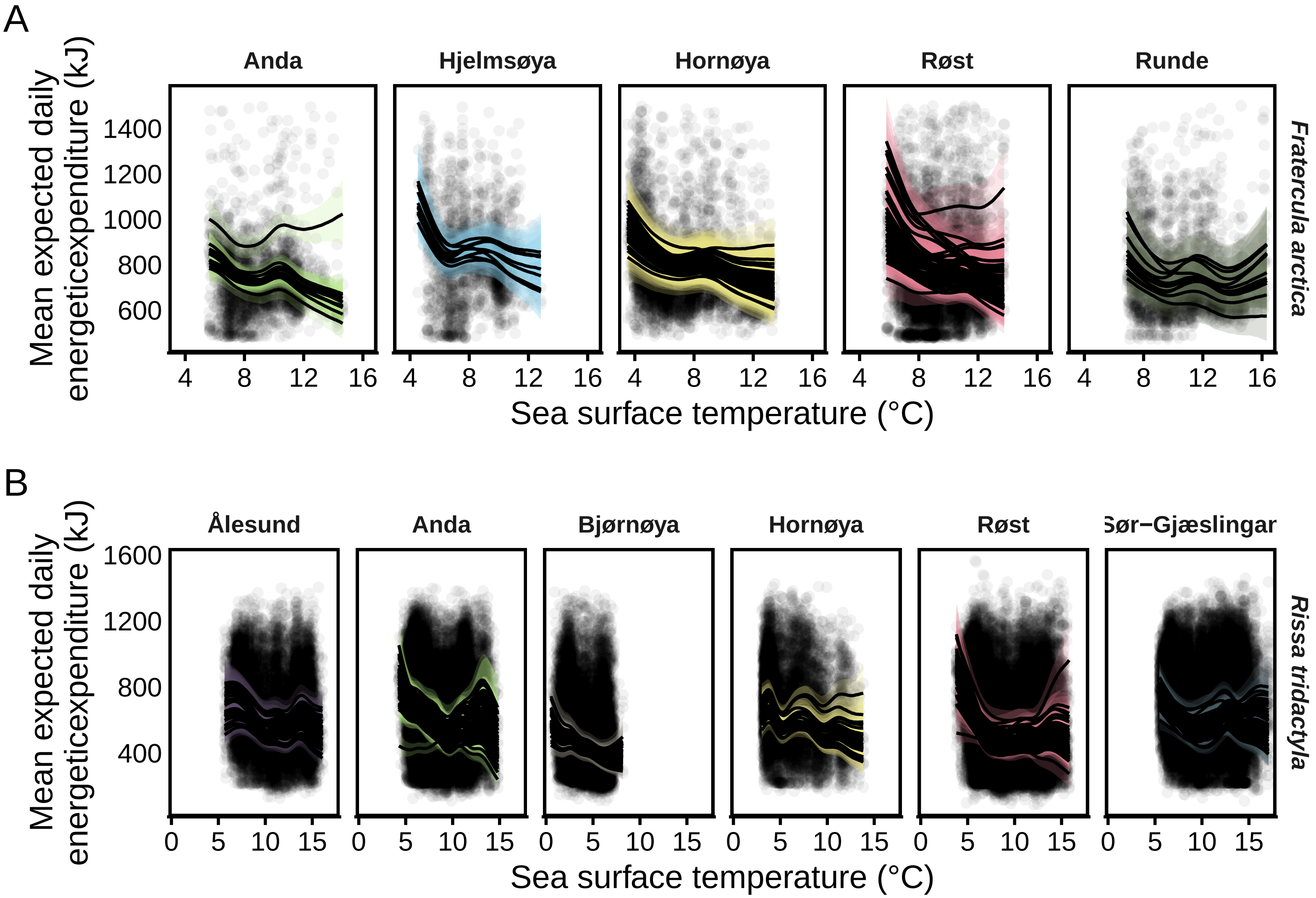


Figure S8: Expected daily energetic expenditure (kJ per day) for each individual and colony, in A) puffins and B) kittiwakes, as predicted by generalised additive models. Solid lines are median estimates, with shaded regions the pointwise 95% credible interval.

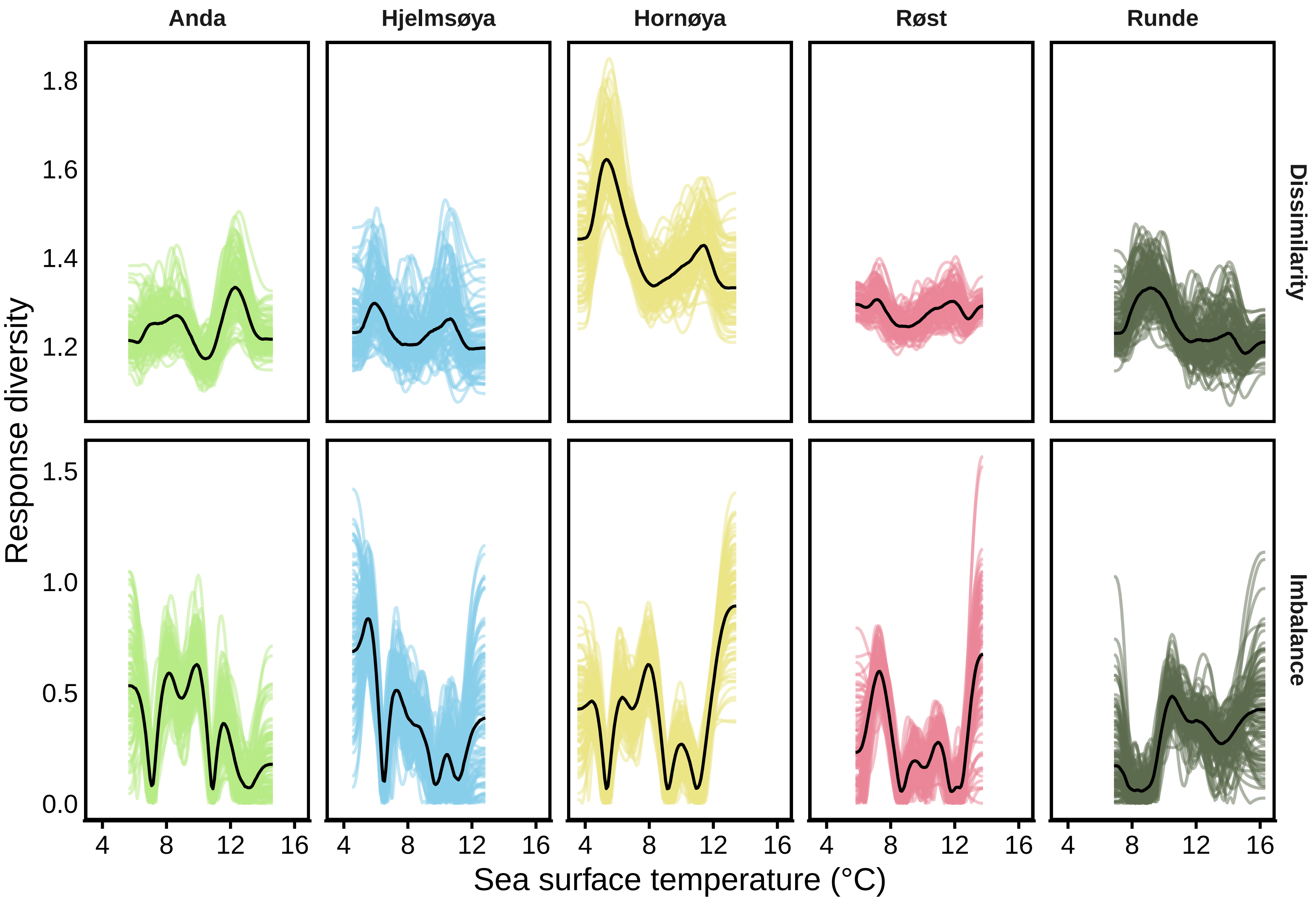


Figure S9: Spaghetti plot of puffin foraging effort response diversity. Solid, dark lines are each colony's median response diversity, while coloured lines are 100 random draws for each.

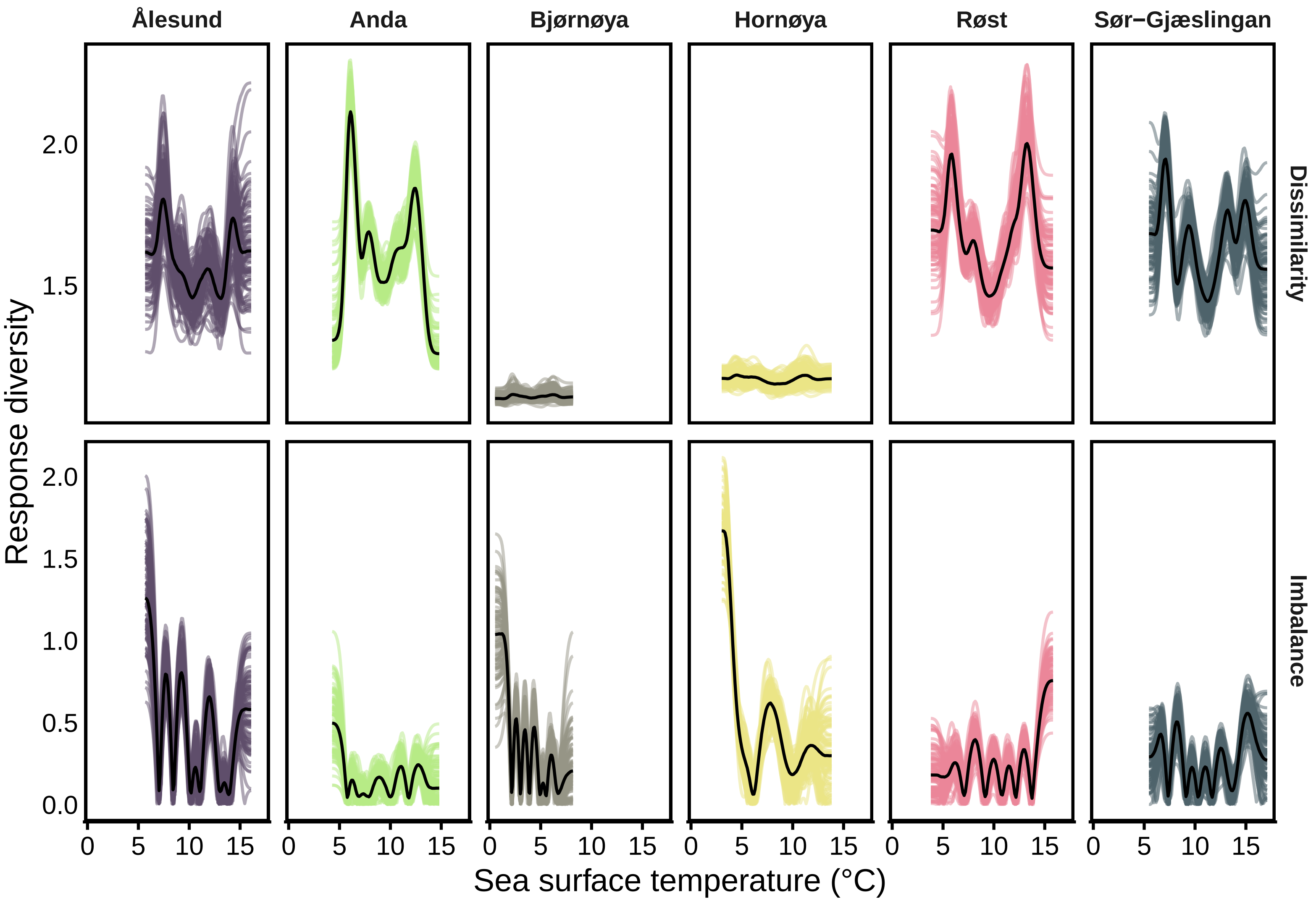


Figure S10: Spaghetti plot of kittiwake foraging effort response diversity. Solid, dark lines are each colony's median response diversity, while coloured lines are 100 random draws for each.

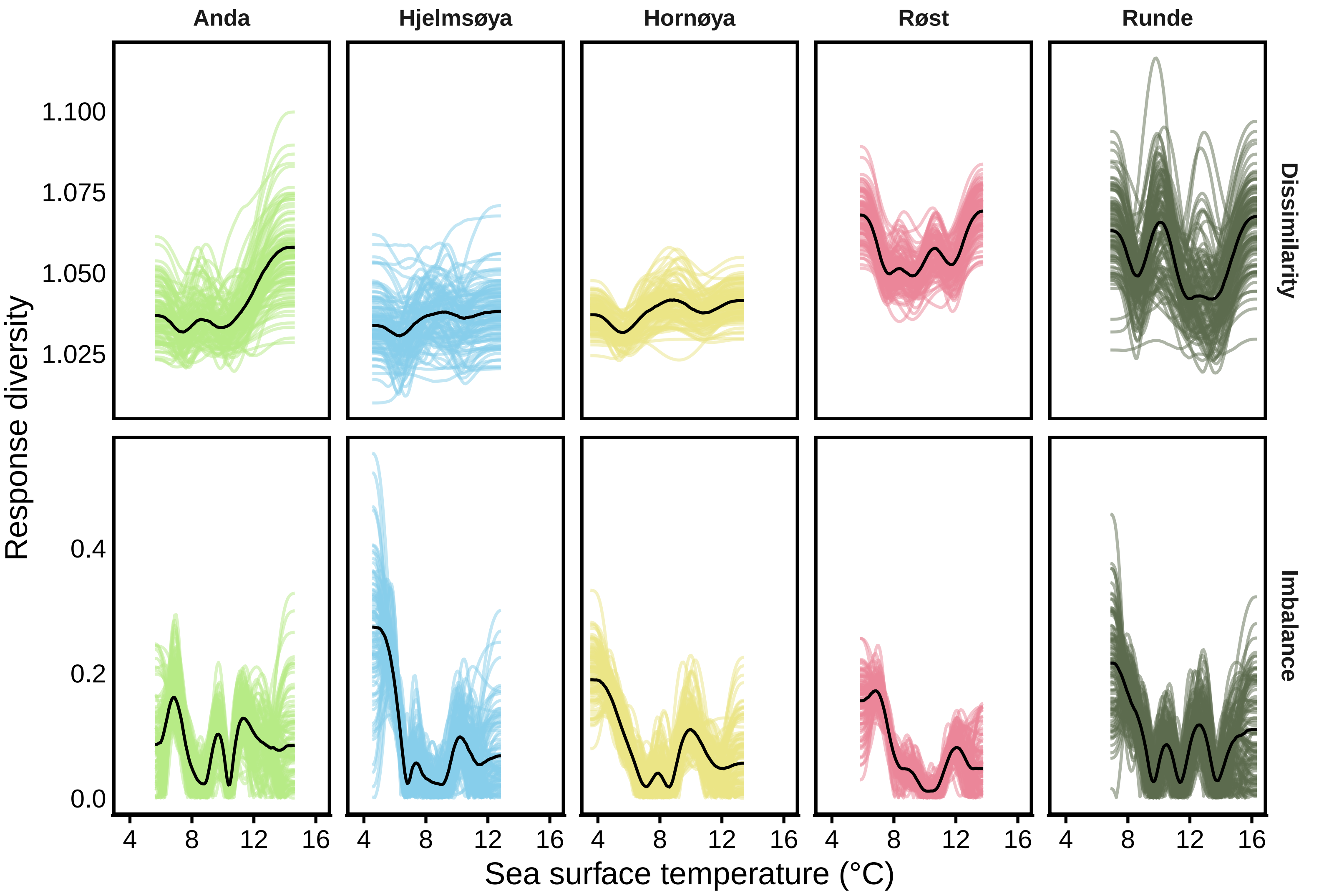


Figure S11: Spaghetti plot of puffin daily energetic expenditure (kJ) response diversity. Solid, dark lines are each colony's median response diversity, while coloured lines are 100 random draws for each.

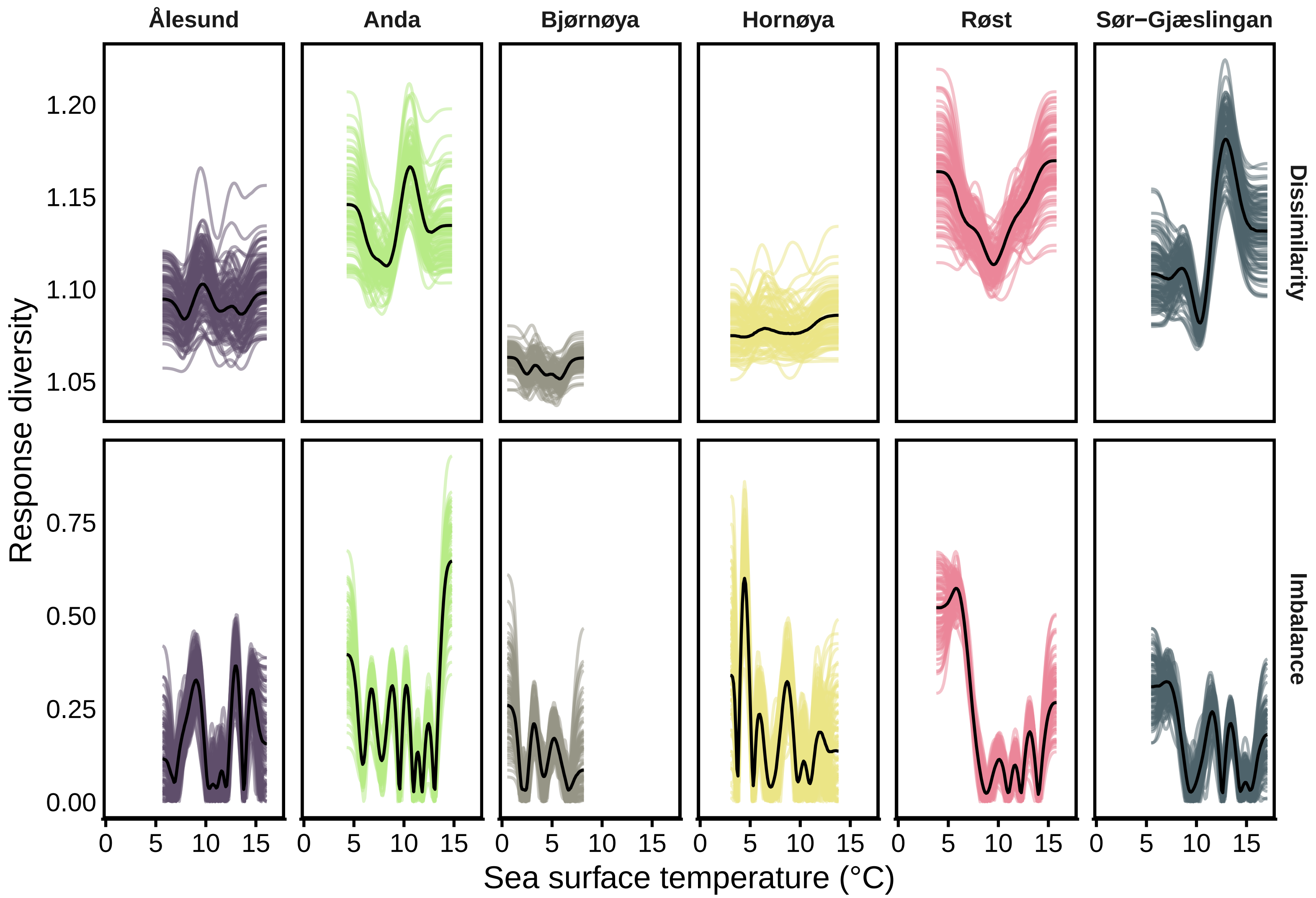


Figure S12: Spaghetti plot of kittiwake daily energetic expenditure (kJ) response diversity. Solid, dark lines are each colony's median response diversity, while coloured lines are 100 random draws for each.

#### S3: Supplementary tables

Table S1: Seabird population measurements and their units.

| Species | Colony | Measurement | Method | Units | Sample size | Coverage |
|---|---|---|---|---|---|---|
| *Fratercula arctica* | Anda | Population counts | Apparently occupied burrows | Number of pairs | 18 yrs | 2005-2023 |
| *Fratercula arctica* | Anda | Behaviour | Global location sensors | Light intensity (lux) & conductivity (proportion of time underwater) | 12 individuals | 2014-2023 |
| *Fratercula arctica* | Hjelmsøya | Population counts | Apparently occupied burrows | Number of pairs | 10 yrs | 2015-2024 |
| *Fratercula arctica* | Hjelmsøya | Behaviour | Global location sensors | Light intensity (lux) & conductivity (proportion of time underwater) | 10 individuals | 2014-2022 |
| *Fratercula arctica* | Hornøya | Population counts | Apparently occupied burrows | Number of pairs | 42 yrs | 1980-2024 |
| *Fratercula arctica* | Hornøya | Behaviour | Global location sensors | Light intensity (lux) & conductivity (proportion of time underwater) | 37 individuals | 2012-2020 |
| *Fratercula arctica* | Røst | Population counts | Apparently occupied burrows | Number of pairs | 42 yrs | 1983-2024 |
| *Fratercula arctica* | Røst | Behaviour | Global location sensors | Light intensity (lux) & conductivity (proportion of time underwater) | 44 individuals | 2014-2023 |
| *Fratercula arctica* | Runde | Population counts | Apparently occupied burrows | Number of pairs | 37 yrs | 1980-2023 |
| *Fratercula arctica* | Runde | Behaviour | Global location sensors | Light intensity (lux) & conductivity (proportion of time underwater) | 11 individuals | 2014-2020 |
| *Rissa tridactyla* | Ålesund | Population counts | Apparently occupied nests | Number of pairs | 13 yrs | 2011-2023 |
| *Rissa tridactyla* | Ålesund | Behaviour | Global location sensors | Light intensity (lux) & conductivity (proportion of time underwater) | 37 individuals | 2015-2022 |
| *Rissa tridactyla* | Anda | Population counts | Apparently occupied nests | Number of pairs | 20 yrs | 2005-2024 |
| *Rissa tridactyla* | Anda | Behaviour | Global location sensors | Light intensity (lux) & conductivity (proportion of time underwater) | 60 individuals | 2009-2023 |
| *Rissa tridactyla* | Bjørnøya | Population counts | Apparently occupied nests | Number of pairs | 35 yrs | 1988-2024 |
| *Rissa tridactyla* | Bjørnøya | Behaviour | Global location sensors | Light intensity (lux) & conductivity (proportion of time underwater) | 38 individuals | 2009-2023 |
| *Rissa tridactyla* | Hornøya | Population counts | Apparently occupied nests | Number of pairs | 43 yrs | 1980-2024 |
| *Rissa tridactyla* | Hornøya | Behaviour | Global location sensors | Light intensity (lux) & conductivity (proportion of time underwater) | 25 individuals | 2012-2021 |

Table S1: Seabird population measurements and their units.

| Species | Colony | Measurement | Method | Units | Sample size | Coverage |
|---|---|---|---|---|---|---|
| *Rissa tridactyla* | Røst | Population counts | Apparently occupied nests | Number of pairs | 37 yrs | 1988-2024 |
| *Rissa tridactyla* | Røst | Behaviour | Global location sensors | Light intensity (lux) & conductivity (proportion of time underwater) | 53 individuals | 2008-2023 |
| *Rissa tridactyla* | Sør-Gjæslingan | Population counts | Apparently occupied nests | Number of pairs | 14 yrs | 2011-2024 |
| *Rissa tridactyla* | Sør-Gjæslingan | Behaviour | Global location sensors | Light intensity (lux) & conductivity (proportion of time underwater) | 53 individuals | 2013-2023 |

Table S2: Pearson correlations estimated between among-individual foraging effort response diversity metrics and population-level stability

| Species | Response diversity metric | Stability metric | Median rho | Lower CI | Upper CI |
|---|---|---|---|---|---|
| Kittiwakes | Dissimilarity | Dominant eigenvalue | -0.869 | -0.940 | -0.769 |
| Kittiwakes | Dissimilarity | Temporal stability | 0.488 | 0.295 | 0.662 |
| Kittiwakes | Imbalance | Dominant eigenvalue | 0.690 | 0.302 | 0.939 |
| Kittiwakes | Imbalance | Temporal stability | -0.383 | -0.773 | 0.081 |
| Puffins | Dissimilarity | Dominant eigenvalue | -0.728 | -0.877 | -0.499 |
| Puffins | Dissimilarity | Temporal stability | -0.309 | -0.608 | 0.046 |
| Puffins | Imbalance | Dominant eigenvalue | 0.322 | -0.232 | 0.784 |
| Puffins | Imbalance | Temporal stability | 0.265 | -0.363 | 0.727 |

Table S3: Pearson correlations estimated between among-individual daily energetic expenditure response diversity metrics and population-level stability

| Species | Response diversity metric | Stability metric | Median rho | Lower CI | Upper CI |
|---|---|---|---|---|---|
| Kittiwakes | Dissimilarity | Dominant eigenvalue | -0.599 | -0.812 | -0.375 |
| Kittiwakes | Dissimilarity | Temporal stability | 0.608 | 0.306 | 0.844 |
| Kittiwakes | Imbalance | Dominant eigenvalue | 0.272 | -0.046 | 0.544 |
| Kittiwakes | Imbalance | Temporal stability | -0.001 | -0.338 | 0.379 |
| Puffins | Dissimilarity | Dominant eigenvalue | 0.144 | -0.400 | 0.613 |
| Puffins | Dissimilarity | Temporal stability | -0.484 | -0.835 | 0.124 |
| Puffins | Imbalance | Dominant eigenvalue | 0.061 | -0.861 | 0.873 |
| Puffins | Imbalance | Temporal stability | -0.318 | -0.927 | 0.760 |
